# Species Delimitation using Genome-Wide SNP Data

**DOI:** 10.1101/001172

**Authors:** Adam D. Leaché, Matthew K. Fujita, Vladimir N. Minin, Remco R. Bouckaert

**Author notes:** Corresponding author: Adam D. Leaché, Department of Biology, 24 Kincaid Hall, University of Washington, Seattle, WA 98195, USA.

## Abstract

The multi-species coalescent has provided important progress for evolutionary inferences, including increasing the statistical rigor and objectivity of comparisons among competing species delimitation models. However, Bayesian species delimitation methods typically require brute force integration over gene trees via Markov chain Monte Carlo (MCMC), which introduces a large computation burden and precludes their application to genomic-scale data. Here we combine a recently introduced dynamic programming algorithm for estimating species trees that bypasses MCMC integration over gene trees with sophisticated methods for estimating marginal likelihoods, needed for Bayesian model selection, to provide a rigorous and computationally tractable technique for genome-wide species delimitation. We provide a critical yet simple correction that brings the likelihoods of different species trees, and more importantly their corresponding marginal likelihoods, to the same common denominator, which enables direct and accurate comparisons of competing species delimitation models using Bayes factors. We test this approach, which we call Bayes factor delimitation (*with genomic data; `BFD*`), using common species delimitation scenarios with computer simulations. Varying the numbers of loci and the number of samples suggest that the approach can distinguish the true model even with few loci and limited samples per species. Misspecification of the prior for population size *θ* has little impact on support for the true model. We apply the approach to West African forest geckos (*Hemidactylus fasciatus* complex) using genome-wide SNP data data. This new Bayesian method for species delimitation builds on a growing trend for objective species delimitation methods with explicit model assumptions that are easily tested.

---

Genomic data are having a dramatic impact on our ability to resolve the tree of life (Faircloth et al. 2012), but delimiting species at the tips of the tree remains to see comparable gains. New species delimitation methods are increasing in statistical rigor and objectivity as a result of adopting a multispecies coalescent model (Fujita et al. 2012), although expanding these methods to truly embrace genome-scale data may be limited by their reliance on gene trees. Individually, gene trees can be estimated quickly using fast heuristic methods (Stamatakis 2006), but combining hundreds or thousands of gene trees into a single species delimitation framework presents serious computational challenges and a poor prognosis for genome-wide species delimitations. Thus far, species delimitation studies using gene trees have been limited to approximately 20 loci (Carstens et al. 2013), but as many studies trend toward large phylogenomic datasets exceeding 100s of loci (O’Neill et al. 2013); (Wagner et al. 2013); (Smith et al. 2013) there is a real need for genomically-enabled species delimitation approaches.

New methods for estimating species trees without gene trees (Bryant et al. 2012); (Patterson et al. 2012) open the doors for a remedy. The `SNAPP` method (Bryant et al. 2012) estimates species trees directly from biallelic markers (e.g., SNP or AFLP data), and bypasses the necessity of having to explicitly integrate or sample the gene trees at each locus. The method works by estimating the probability of allele frequency change across ancestor/descendent nodes. Given a species tree, the probability of the allele frequencies at a given locus is the probability of a site given a gene tree multiplied by the probability of the gene tree given the species tree, summed over all possible gene tree topologies and integrated over all possible gene tree branch lengths (Bryant et al. 2012). The result is a posterior distribution for the species tree, species divergence times, and effective population sizes, all obtained without the estimation of gene trees.

Comparisons among candidate species delimitation models that contain different numbers of species is relatively easy with the use of Bayes factors (Grummer et al. 2013), and the existing approach (Bayes factor delimitation; `BFD`) uses traditional DNA sequence data to simultaneously estimate gene trees and species trees. The approach requires the estimation of marginal likelihoods for each competing model, and recent work provides a number of solutions for estimating these values (Baele et al. 2012). Bayes factor species delimitation has a number of advantages over other Bayesian species delimitation approaches (Yang and Rannala 2010). A significant advantage is the ability to integrate over species trees during the species delimitation procedure, which removes the constraint of specifying a guide tree that represents the true species relationships. The species tree is usually uncertain, and incorrect guide trees can bias Bayesian species delimitation (Leaché and Fujita 2010). Another advantage is the ability to compare non-nested models that contain different numbers of species, or different assignments of samples to species (Grummer et al. 2013). Currently, the approach has several limitation. It’s use for heuristically searching among all possible species assignments is hampered by the need to predefine the number of species and sample assignments. Methods including the `GMYC` (Pons et al. 2006), `SpeDeSTEM` (Ence and Carstens 2011), and `BROWNIE` (O’Meara 2010) are more appropriate heuristic tools for producing species assignments when the researcher has no preconceived species delimitation models to test. Finally, `BFD`, as well as the other species delimitation methods listed previously, rely at some point on gene tree estimation, and are therefore not easily extended to genome data.

Here, we incorporate Bayesian multispecies coalescent species delimitation using genome-wide SNP data (or other types of biallelic markers, including AFLP data into the `SNAPP` method), but with a critical new addition. First, we describe the approach and show that marginal likelihoods for alternative species delimitation models are not directly comparable. We solve this problem by adding proportionality constants that bring marginal likelihoods to the same scale for comparing competing species trees with Bayes factors. Second, we use computer simulations to verify that our approach works over broad parameter and data quantity values when the number of species is known. Finally, we conduct empirical species delimitation using genome-wide SNP data for West African forest geckos (*Hemidactylus fasciatus* complex). We test a four-species hypothesis for this group that was supported by a previous study (Leaché and Fujita 2010) based on five independent nuclear loci and Bayesian species delimitation (`BPP`).

## Species Delimitation Without Gene Trees

We start with *n* individuals and *m* unlinked biallelic markers typed in these individuals, with alleles designated as 0 and 1. Suppose each individual is assigned to one of *k* populations/species. A Bayesian method implemented in the software package `SNAPP` (Bryant et al. 2012) uses the marker data and species assignments to estimate the species phylogeny. However, the assignment of individuals to species and the number of species is often uncertain, so we would like to compare multiple species delimitations that make different assumptions about sample assignments and *k*. Such comparisons are straightforward to accomplish in a Bayesian framework using Bayes factors (Kass and Raftery 1995). Suppose we want to use an *n* × *m* matrix of biallelic markers **y** to compare two competing species assignments **a**_1_ = (*a*_11_,…,*a*_1*n*_) and **a**_2_ = (*a*_21_,…,*a*_2*n*_), where *a*_1*j*_ and *a*_2*j*_ are the corresponding species indicators for individual *j*. For example, an assignment **a** = (1, 1, 2, 2, 3, 3, 1) says that individuals 1, 2, 7 belong to species 1, individuals 3 and 4 belong to species 2, and individuals 5 and 6 belong to species 3. The Bayes factor comparing the two assignments is

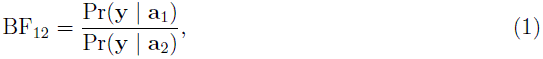

where

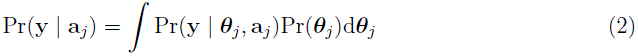

is a marginal likelihood, ***θ**_j_* is a set of all model parameters that define a species tree model corresponding to species assignment **a***_j_*, Pr(**y** *| **θ**_j_*, **a***_j_*) is the likelihood function, and Pr(***θ**_j_*) is a prior density of model parameters (we will use Pr(*·*) to designate both probability and density). In order to compare multiple species delimitation models, one can rank these models by their corresponding marginal likelihoods (2). Selecting the highest ranked model is a statistically consistent procedure, meaning that the highest ranked model is guaranteed to be the correct model as the amount of data increases, assuming the correct model is being considered. The software package `SNAPP` has an implementation of path sampling to approximate marginal likelihoods of the form (2), but the current implementation of the software rescales these likelihoods in a way that makes marginal likelihoods incomparable. We explain this rescaling and the corresponding remedy below.

## Proportionality Constants for Bayes Factor Species Delimitation

We use `SNAPP` (Bryant et al. 2012); (RoyChoudhury et al. 2008) to bypass the explicit integration over the space of gene trees using an algorithm for computing the likelihood

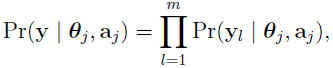

where *l* indexes one of *m* unlinked loci under study, **y***_l_* is a vector of markers at locus *l* and **y =** (**y**_1_,…,**y***_m_*). The above algorithm starts by compressing the data matrix **y** into sufficient statistics. For each locus *l*, sufficient statistics for species tree estimation are **s***_l_* = (*s_l_*_1_,…,*s_lk_*) and **n***_l_* = (*n_l_*_1_,…,*n_lk_*), where *s_li_* is the number of 1 alleles at locus *l* in individuals belonging to species *i* and *n_li_* is the number of individuals with nonmissing data at locus *l* in species *i*, and *k* is the number of species (Bryant et al. 2012); (RoyChoudhury et al. 2008). Therefore, associated with these sufficient statistics is the likelihood

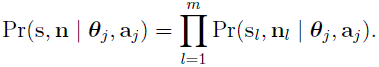

As a result, the path sampling implemented in `SNAPP` computes the marginal likelihood,

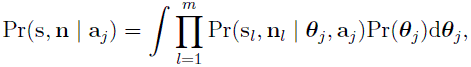

that also corresponds to the sufficient statistics rather than to the original data. This procedure does not affect the estimation of the species tree, the original objective of `SNAPP`, but since different species assignments change the allele counts within species (i.e. the sufficient statistics), the marginal likelihoods provided by `SNAPP` for different species delimitation models are incomparable. However, the original marker data likelihood is equal to the sufficient statistics data likelihood up to a proportionality constant. By computing this constant for each species assignment, we can bring `SNAPP` marginal likelihoods to the same scale and perform model ranking and Bayes factor calculations.

Let 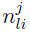 be the number of individuals with nonmissing data at locus *l* in species *i* under species assignment **a***_j_* and let 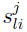 be the number of 1 alleles at locus *l* in individuals belonging to species *i* under species assignment **a***_j_*. Notice that the number of species *k_j_* depends on *j*. Then, for locus *l* and assignment *j*,

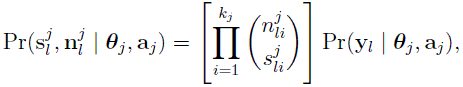

where 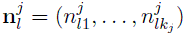 and 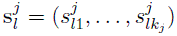. This means that we can compute the corrected marginal likelihood of species assignment **a***_j_* as

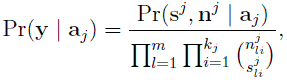

where 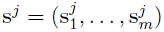, 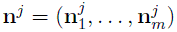 and 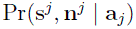 is the marginal likelihood that can be approximated by `SNAPP`. Marginal likelihoods of two species delimitation models can be plugged into formula (1) to compare the models using Bayes factors.

## Materials and Methods

### Computer Simulations

The multispecies coalescent SNP simulator `SimSNAPP` (Bryant et al. 2012) was used to generate polymorphic biallelic markers on a predefined species tree (100 replicates per simulation). The species tree is asymmetric and contains four species (Fig. 1), and is equivalent in divergence times and population sizes to a species tree used in two other simulation studies (Liu and Pearl 2007); (Bryant et al. 2012). We alter the species assignments of the true model to test several common species delimitation scenarios, including species lumping (two species are combined into one), splitting (one species is arbitrarily split into two species), and misclassification of samples to different species. We simulated SNP datasets containing either 100, 500, or 1000 polymorphic biallelic characters. Each separate marker is unlinked given the species tree. We conducted additional simulations sampling either two, five, or ten samples per species to examine the influence of sampling design on species delimitation.

**Figure 1:**
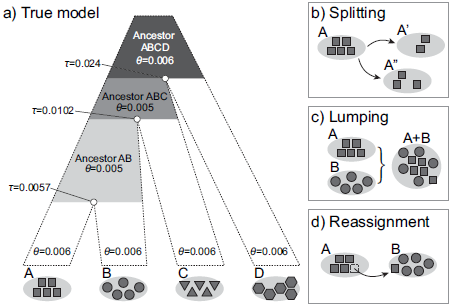
The fully-specified species tree used to simulate SNP data for Bayes factor species delimitation (a). Perturbations to the true model include (b) splitting a species into two false species, (c) lumping two distinct species into one, and (d) reassigning a sample into the wrong species. Simulations are conducted with SNP matrices of different sizes (100, 500, 1000), variable sampling within species (2, 5, 10), and with different theta priors (correct, high, low). Species tree divergence times are in units of expected mutations per site.

Our simulation framework allows us to properly specify the prior distributions for parameters that most empiricists typically must estimate from the data. In most cases, we specified prior distributions to closely match the values used during simulation. Doing so allowed us to focus on the performance of the species delimitation framework instead of problems associated with prior misspecification. We analyzed the simulated data using a correctly specific prior for the population size parameter, *θ* (gamma(2,333)), which results in an accurate prior mean = 0.006. However, we are also interested in understanding how misspecified priors might impact our ability to accurately delimit species. Therefore, we also conducted simulations with a gamma prior off target by orders of magnitude to result in a “low” prior mean = 0.0001 (gamma(2,20000)), and a “high” prior mean = 0.01 (gamma(2,200)).

We analyzed the simulated datasets using a modified version of `SNAPP` that includes proportionality constants. Posterior probability distributions for the species tree, divergence times, and population sizes are a product of the analysis, yet estimating the marginal likelihood is the immediate goal for Bayes factor model comparison. Estimating the marginal likelihood requires extra computation compared to typical Bayesian inference (Baele et al. 2012), and path sampling and stepping-stone methods work well in the context of Bayes factor delimitation of species using multilocus DNA sequence data (Grummer et al. 2013). We conducted path sampling with 48 steps to estimate the marginal likelihood. Each path sampling step is a power posterior differing only in its power, *β*. When *β* = 1 the samples are taken from the posterior and the analysis is informed by the data. When *β* = 0, the prior is sampled without any influence of the data (Baele et al. 2012). Intermediate *β* values alter the ratio of data and prior, and therefore path sampling can accurately estimate the marginal likelihood (Baele et al. 2012). An Markov chain Monte Carlo (MCMC) chain length of 200000 with a pre-burnin of 50000 was sufficient to achieve large effective sample sizes and apparent stationarity.

The strength of support from Bayes factor (BF; equation 1) comparisons of competing models was evaluated using the framework of (Kass and Raftery 1995). A positive BF test statistic (2 × log_*e*_) reflects evidence in favor of model 1, whereas negative BF values are considered as evidence favoring model 2. The BF scale is as follows: 0 < 2 × log_*e*_ BF < 2 is not worth more than a bare mention, 2 < 2 × log_*e*_ BF > 6 is positive evidence, 6 < 2 × log_*e*_ BF < 10 is strong support, and 2 × log_*e*_ BF > 10 is decisive.

### Empirical Data

We applied `BFD*` to new SNP data collected for West African forest geckos in the *Hemidactylus fasciatus* complex. A previous species delimitation study utilizing five nuclear loci found strong support for at least four unique evolutionary lineages within *Hemidactylus fasciatus* using the Bayesian species delimitation method `BPP` (Leaché and Fujita 2010). The validity of the four species was debated (Bauer et al. 2011); (Fujita and Leaché 2011), but we consider the four species scenario (Fig. 2a) a logical starting point for testing competing species delimitation models (Fig. 2b–g). The alternative species delimitation models that we test include lumping species (Fig. 2b–d), splitting species (Fig. 2e–f), and reassigning samples between species (Fig. 2g).

**Figure 2:**
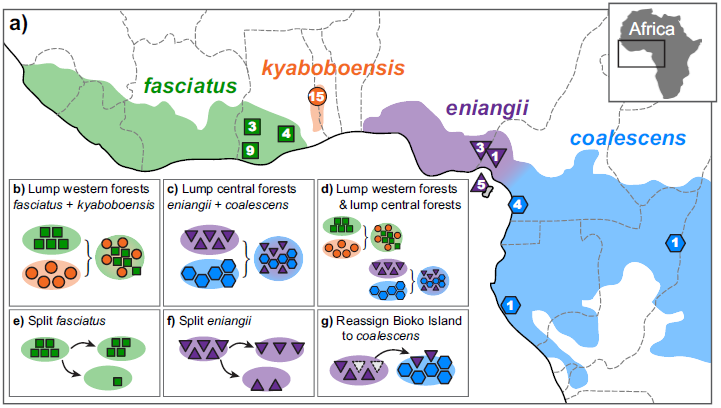
Geographic sampling of *Hemidactylus fasciatus* complex geckos (numbers in symbols indicate sample sizes), and our preferred current taxonomy (a). BFD* is used to test the alternative species delimitation models outlined in b-g.

To collect genome-wide sampling of SNPs, we followed the laboratory protocols for double-digest RADseq (ddRADseq) described in (Peterson et al. 2012). We collected data for 46 West African forest geckos in the *Hemidactylus fasciatus* complex, allocated into our preferred taxonomy as follows: *H. coalescens* (n = 6; Cameroon, Congo, Gabon), *H. eniangii* (n = 9; Cameroon, Equatorial Guinea, Nigeria), *H. fasciatus* (n = 16; western Ghana), and *H. kyaboboensis* (n = 15; Ghanas Togo Hills). By simultaneously characterizing and genotyping samples, ddRADseq has the desirable property of greatly reducing ascertainment bias (Hohenlohe et al. 2010); (Helyar et al. 2011). For each individual, we extracted high-molecular weight genomic DNA from liver or muscle tissue, checked the quality on agarose gels and measured the concentration using a Qubit. Overnight double digestion with SbfI and MspI used 0.5 μg of DNA. Fragments were purified with Agencourt AMPure beads before ligation of barcoded Illumina adaptors onto the fragments. Equimolar amounts of each sample were pooled prior to size selection using a Blue Pippin Prep. Final library amplification used proofreading Taq and Illumina’s indexed primers. We used two quality control measures prior to sequencing, including quantitative PCR to accurately measure DNA concentration of adaptor-associated fragments, and a BioAnalyzer run to confirm the sizes of fragments. The final libraries were sequenced (50-bp, single-end run) on one Illumina HiSeq 2000 lane at the Vincent J. Coates QB3 Genomic Sequencing Facility at UC Berkeley.

Raw Illumina reads were filtered for contaminating adaptors and primers using the `FASTX-Toolkit`. We processed the filtered data using `STACKS` (Catchen et al. 2011), a group of programs and scripts that perform additional filtering based on sequence quality and identifies putative loci and haplotypes for each individual, and organizes them into a MySQL database. Working per individual, we used ustacks to create putative loci by grouping reads that differ by a threshold of three mismatches. This threshold amounts to 7.7% nuclear divergence for 39-bp fragments (50-bp sequence minus the five-bp barcode and partial SbfI site), which we view as high for intraspecific diversity but necessary to capture potential admixture between divergent populations. The program ustacks then uses a maximum likelihood algorithm to determine haplotypes for each individual (Hohenlohe et al. 2010). We removed putative loci with more than twice the standard deviation of coverage depth to filter out repetitive elements and stacks of paralogous loci. Unique loci from all individuals were aggregated into a “Catalog” using cstacks, keeping track of the haplotype diversity within each locus with a mismatch threshold of three, which reflects a range of divergences that could constitute potential cryptic species. We resolved haplotypes for each individual for each locus in the catalog using sstacks. The program populations outputs haplotype files from which we reconstituted alignments for downstream analyses using our own scripts. Since each RAD locus may contain multiple linked SNPs, we assembled a final data matrix containing a single SNP selected at random from each locus.

We analyzed two assemblies of the empirical data that differed in the level of missing data. One assembly contained no missing data, and therefore had only a small number of loci (57), whereas lowering our tolerance for missing data to 10% resulted in a matrix containing 1202 loci. We analyzed these data using the modified version of SNAPP, which was implemented as a plug-in to the BEAST 2 (Bouckaert et al. 2013). We conducted path sampling with 48 steps (100000 MCMC steps, 10000 pre-burnin steps) to estimate the marginal likelihood. The software is open source and is available for download from http://code.google.com/p/snap-mcmc/. We have created a wiki-page for BFD* that provides detailed steps on how to install the program, set up the XML file, and run the analyses (http://www.beast2.org/wiki/index.php/BFD*/).

## Results

### The Importance of Proportionality Constants in Species Tree Comparisons

A comparison of marginal likelihood values estimated both with and without proportionality constants, and their influence on Bayes factor comparisons of species trees, is shown in Figure 3. Under simulation, uncorrected marginal likelihoods are higher compared to their corrected counterparts estimated with proportionality constants (Fig 3a). Arbitrarily lumping species reduces the number of parameters, yet the uncorrected marginal likelihood values increase, which is a counterintuitive result that is rectified with the use of corrected marginal likelihoods. The incorrect estimation of marginal likelihoods leads to a strong bias in BF comparisons of species trees, and the uncorrected values tend to reject the true model in favor of less complex models containing fewer species (i.e., models that lump species) with decisive support (Fig. 3b).

**Figure 3:**
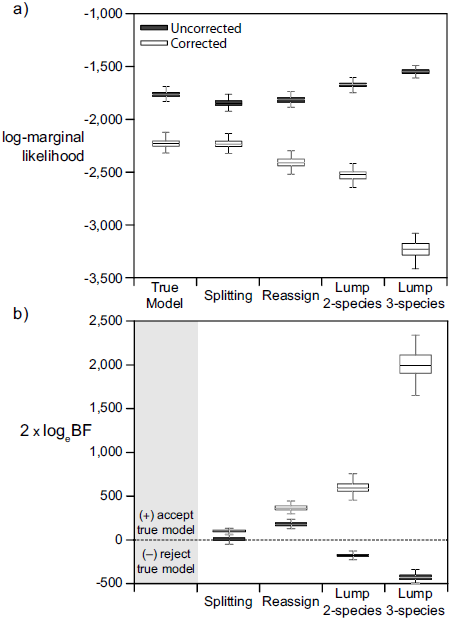
Comparisons of the behavior of corrected and uncorrected marginal likelihoods (a), and their influence on Bayes factor comparisons of candidate species trees (b). The simulated data used in this comparison include 500 SNPs and 5 samples per species.

Bayes factor comparisons of candidate species trees with corrected marginal likelihoods consistently favor the true model over the competing models as reflected by the positive BF values (Table 1). Reassigning samples to the wrong species, or lumping species together are rejected decisively across all simulations (2 × log_*e*_ BF ≥ 70; Table 1). Arbitrarily splitting a species into two putative sister species produces strong 2 × log_*e*_ BF values (>6), and although most simulation replicates supported the true model, some replicates supported the alternative model when the *θ* prior was misspecified (Table 1). This result indicates that species delimitation models that incorrectly parse weakly diverged populations, or that split populations connected by moderate to high gene flow, could be difficult to distinguish using BFD*. Acquiring decisive BF support for these spurious grouping may be quite difficult.

**Table 1:** Simulation results for BFD* species delimitation. Results are mean values (and standard deviations) across 100 simulation replicates.

| Loci | Samples | $\theta$ prior | Marginal likelihood ($\log_e$) | | | | Bayes factor ($2 \times \log_e$) | | |
| --- | --- | --- | --- | --- | --- | --- | --- | --- | --- |
|  |  |  | true | reassign | lump | split | reassign | lump | split |
| 500 | 5 | correct | -2228.3 (45.8) | -2411.2 (53.3) | -2527.4 (58.7) | -2234.0 (50.0) | +365.8 (41.2) | +598.2 (61.8) | +8.4 (1.4) |
| 500 | 5 | low | -2224.4 (45.9) | -2411.1 (54.1) | -2531.8 (60.2) | -2228.6 (46.0) | +373.5 (42.9) | +614.9 (64.8) | +8.5 (1.5) |
| 500 | 5 | high | -2229.8 (38.0) | -2411.4 (43.9) | -2527.2 (47.2) | -2234.1 (39.0) | +363.2 (33.0) | +594.8 (62.1) | +8.5 (17.2) |
| 100 | 5 | correct | -449.7 (20.5) | -486.3 (23.0) | -508.0 (24.9) | -452.9 (20.4) | +73.3 (20.9) | +116.6 (29.7) | +6.4 (1.8) |
| 1000 | 5 | correct | -4441.1 (73.8) | -4797.8 (87.1) | -5028.2 (98.5) | -4445.9 (73.7) | +713.3 (72.0) | +1174.2 (108.4) | +9.5 (1.6) |
| 500 | 2 | correct | -1072.2 (35.3) | -1139.1 (40.7) | -1136.1 (40.7) | -1075.6 (35.2) | +133.8 (27.2) | +127.8 (27.1) | +6.8 (1.2) |
| 500 | 10 | correct | -3143.3 (81.4) | -3358.0 (82.8) | -3746.7 (105.6) | -3148.2 (81.8) | +429.3 (59.3) | +1206.8 (148.6) | +9.8 (1.7) |

### Sampling Intensity

We might expect that fluctuations in sampling intensity (i.e., loci or individuals) might impact our ability to discriminate alternative species delimitation models. Species tree simulation studies have demonstrated that accuracy increases with the number of loci (Bryant et al. 2012); (Heled and Drummond 2010); (Leaché and Rannala 2011), and we expect that discriminating the true species delimitation model from alternatives should become easier with more data. Under the simulation conditions used here, species delimitation with as few as 100 SNPs is adequate for at least strong (split model) or decisive (reassignment and lump models) BF support for the true model (Table 1). Increasing the number of SNPs results in more decisive BF support for the true model. However, BF support for the true model in relation to the split model provides only marginal improvement with even up to 1000 SNPs (Table 1).

Coalescent methods can also gain information by including more individuals for each species (Maddison and Knowles 2006). We find that increasing the number of samples for each species increases the BF support for the true model (Table 1). This pattern is strongest under models that reassign samples or lump species, which become easier to distinguish from the true model as the number of samples increases (while holding the number of loci constant). For instance, lumping species produces an average 2 × log_*e*_ BF score of +27.1 when including only two samples per species, but the 2 × log_*e*_ BF score increases to +61.8 and +148.6 when sampling five or 10 samples per species, respectively (Table 1). Arbitrarily splitting a species is the most difficult scenario to distinguish from the true model, and adding more samples per species adds a relatively small contribution to the BF scores. The 2 × log_*e*_ BF scores for two, five, and 10 species under the split model are +6.8, + 8.4, and + 9.8, respectively (Table 1).

### Empirical Data

Sequencing on the Illumina HiSeq 2000 platform provided 124+ million raw sequence reads, and resulted in an average SNP coverage of 127.6x. We find the typical trade-off between missing data and high sample representation as is found in other studies utilizing RADseq data (Wagner et al. 2013); (Rubin et al. 2012); (Cariou et al. 2013), namely, the data matrix with the lowest tolerance for missing data has the fewest number of SNPs shared across all samples, and the data matrix with the largest numbers of SNPs has the largest proportion of missing data. For example, a data matrix that is 99.2% complete contains only 129 SNPs, whereas a data matrix allowing 7% missing data (i.e., a SNP can be missing in ≤3 of the 46 samples) contains 1087 SNPs.

Our preferred four-species model is rejected in favor of a five-species model that splits the Bioko Island samples of *H. eniganii* into a separate species, and this result receives decisive support using a small data matrix with 129 SNPs and no missing data, as well as a larger matrix of 1087 SNPs with approximately 10% missing data. Models that lump species or reassign samples between species consistently rank low and are rejected decisively using Bayes factors (Table 2). The specific ranking for these alternative models fluctuates with the number of SNPs, which indicates that either the amount of information content and/or missing data have the potential to impact model ranks. Ultimately, the specific rankings for these models seem unimportant given their decisive rejection.

**Table 2:**
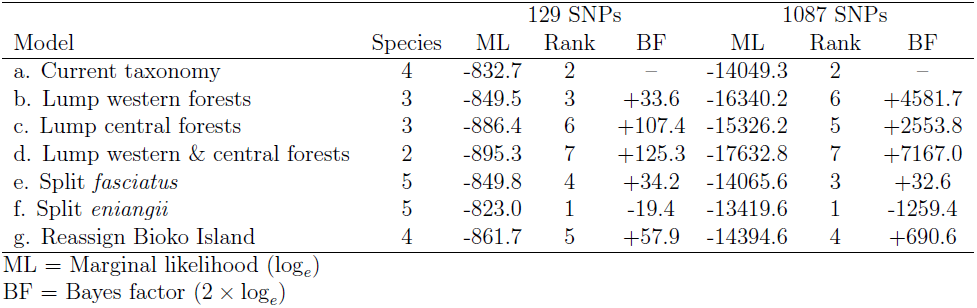
Empirical results for BFD* species delimitation in the *Hemidactylus fasciatus* complex. Models are presented in Figure 3.

| Model | Species | 129 SNPs |  |  | 1087 SNPs |  |  |
| --- | --- | --- | --- | --- | --- | --- | --- |
|  |  | ML | Rank | BF | ML | Rank | BF |
| a. Current taxonomy | 4 | -832.7 | 2 | – | -14049.3 | 2 | – |
| b. Lump western forests | 3 | -849.5 | 3 | +33.6 | -16340.2 | 6 | +4581.7 |
| c. Lump central forests | 3 | -886.4 | 6 | +107.4 | -15326.2 | 5 | +2553.8 |
| d. Lump western & central forests | 2 | -895.3 | 7 | +125.3 | -17632.8 | 7 | +7167.0 |
| e. Split *fasciatus* | 5 | -849.8 | 4 | +34.2 | -14065.6 | 3 | +32.6 |
| f. Split *eniangii* | 5 | -823.0 | 1 | -19.4 | -13419.6 | 1 | -1259.4 |
| g. Reassign Bioko Island | 4 | -861.7 | 5 | +57.9 | -14394.6 | 4 | +690.6 |

ML = Marginal likelihood ( $\log_e$ )
BF = Bayes factor ( $2 \times \log_e$ )

## Discussion

### Hemidactylus Species Delimitation

Allopatric divergence is the primary mechanism producing diversity in geckos belonging to the *Hemidactylus fasciatus* complex. These geckos are restricted to rainforest habitats, and their distributions match those of the major blocks of rainforest. Four species in the group have become reproductively isolated from each other as a result of habitat fragmentation, which has driven allopatric speciation, and one new species appears to be the result of island colonization. The Bioko Island in the Gulf of Guinea harbors a distinct species that is not currently described. We previously found significant support for recognizing the Bioko Island population as a distinct species using five nuclear genes, though we took a“conservative” approach and only recognized four species (Leaché and Fujita 2010). The new SNP data analyzed here provide decisive Bayes factor support for a five-species model that allocates the Bioko Island samples into a separate species (Table 2).

There is a growing suite of coalescent-based species delimitation methods that instill greater objectivity into species delimitation when the appropriate genetic data are available. Understandably, these methods have set afire the necessary discussions over the merit and utility of coalescent-based species delimitation, particularly with concerns over the bias in the rate of lumping or splitting. With the statistical framework of coalescent-based species delimitation methods, these failure to split or lump largely fall into the categories of type I and type II error rates. Common to all methods of species delimitation is the ultimate goal of accurately documenting and quantifying biodiversity that can provide a stable taxonomy. Because of their greater transparency and objectivity, coalescent-based species delimitation methods are an important step forward in attaining this goal.

### Genomic Species Delimitation

Combining data from morphology, ecology, behavior, and genetics is the goal of a pluralistic integrative taxonomy (Padial et al. 2010); (Leaché et al. 2009). While there is currently no inferential framework that can analyze these disparate data types jointly, a component of growing importance is the development of more objective species delimitation tools that take advantage of coalescent theory (Fujita et al. 2012). Indeed, recent progress in statistical species delimitation has largely focused on genetic data (Fujita et al. 2012); (Carstens et al. 2013), likely due to the ease with which they can be abundantly collected even for non-model organisms. Species delimitation with genomic data has desirable properties for systematics, including well-established methodological and statistical foundations, more transparent objectivity over other datatypes, such as morphology (Fujita et al. 2012), and easily-tested model assumptions pertaining to gene flow, selection, or population substructure. Currently, RADseq and sequence capture approaches allow the collection of thousands of loci for hundreds of individuals, producing datasets that are too large and complex for traditional Bayesian and maximum likelihood analyses; indeed, the accumulation of data is outpacing the development of appropriate analytical tools (Sousa and Hey 2013). Nevertheless, we have developed a phylogenomic approach that utilizes SNPs—an important source of genetic variation—to accomplish species delimitation and the documentation of biodiversity, a central goal of systematics with large ramifications for all of biology (Fujita et al. 2012).

### Marginal Likelihoods and Bayes Factors

Obtaining accurate estimates for the marginal likelihood of models forms the foundation for Bayes factor model comparison. The harmonic mean estimator consistently overestimates the marginal likelihood, and is the least desirable approach (Baele et al. 2012). The power posterior approaches including path sampling and stepping-stone sampling both work well in the context of species delimitation (Grummer et al. 2013). Estimating the marginal likelihood using power posteriors requires a substantial amount of computational effort. For example, analyses of our 1202 locus empirical dataset using path sampling required approximately 14.6 days of computation time on an Intel Xeon E5-2650 2.0 GHz 16 core computer with 32 GB of memory. These computation times are not insurmountable when investigating a small set of candidate models, but they could become untenable if the approach is used to rank a large number of models. New methods for Bayesian model selection, including “model-switch stepping-stone sampling”, can decrease the computation time by directly estimating the Bayes factor between two competing models instead of estimating the marginal likelihood for each model separately (Baele et al. 2013). Alternatively, one could try to approximate the joint posterior distribution of species trees and species assignments via MCMC (Choi and Hey 2011), but scaling this approach to genomic data will be challenging computationally.

Using Bayes factor delimitation of species overcomes some of the pitfalls of alternative species delimitation approaches, perhaps the most obvious being circumventing the need to predefine a fixed species tree, which can result in biased support for incorrect models (Leaché and Fujita 2010). This is accomplished by integrating over species trees and other model parameters during marginal likelihood calculations. In addition, marginal likelihoods provide a convenient way to rank alternative, even non-nested, models with the advantage of automatic model complexity penalization (Baele et al. 2012). Finally, marginal likelihood ranking and Bayes factor model comparison do not require the taxonomist to assign prior probabilities to alternative models, which seem difficult to specify for the case of species delimitation.

## Supplementary Material

Supplementary material, including data files and/or online-only appendices, can be found in the Dryad data repository at http://datadryad.org, doi:xx.xxx/dryad.xxxxx.

## Acknowledgments

We thank the University of Washington eSciences Institute for providing computing infrastructure. We thank J. Felsenstein’s Population Genetics reading group for their comments, and X anonymous reviewers for helpful comments.

